# A facile Q-RT-PCR assay for monitoring SARS-CoV-2 growth in cell culture

**DOI:** 10.1101/2020.06.26.174698

**Authors:** Christian Shema Mugisha, Hung R. Vuong, Maritza Puray-Chavez, Sebla B. Kutluay

## Abstract

Severe acute respiratory syndrome coronavirus 2 (SARS-CoV-2), the etiological agent of the ongoing COVID-19 pandemic, has infected millions within just a few months and is continuing to spread around the globe causing immense respiratory disease and mortality. Assays to monitor SARS-CoV-2 growth depend on time-consuming and costly RNA extraction steps, hampering progress in basic research and drug development efforts. Here we developed a facile Q-RT-PCR assay that bypasses viral RNA extraction steps and can monitor SARS-CoV-2 replication kinetics from a small amount of cell culture supernatants. Using this assay, we screened the activities of a number of entry, SARS-CoV-2- and HIV-1-specific inhibitors in a proof of concept study. In line with previous studies which has shown that processing of the viral Spike protein by cellular proteases and endosomal fusion are required for entry, we found that E64D and apilimod potently decreased the amount of SARS-CoV-2 RNA in cell culture supernatants with minimal cytotoxicity. Surprisingly, we found that macropinocytosis inhibitor EIPA similarly decreased viral RNA in supernatants suggesting that entry may additionally be mediated by an alternative pathway. HIV-1-specific inhibitors nevirapine (an NNRTI), amprenavir (a protease inhibitor), and ALLINI-2 (an allosteric integrase inhibitor) modestly inhibited SARS-CoV-2 replication, albeit the IC_50_ values were much higher than that required for HIV-1. Taken together, this facile assay will undoubtedly expedite basic SARS-CoV-2 research, be amenable to mid-throughput screens to identify chemical inhibitors of SARS-CoV-2, and be applicable to a broad number of RNA and DNA viruses.

**Importance:** Severe acute respiratory syndrome coronavirus 2 (SARS-CoV-2), the etiological agent of the COVID-19 pandemic, has quickly become a major global health problem causing immense respiratory disease and social and economic disruptions. Conventional assays that monitor SARS-CoV-2 growth in cell culture rely on costly and time-consuming RNA extraction procedures, hampering progress in basic SARS-CoV-2 research and development of effective therapeutics. Here we developed a facile Q-RT-PCR assay to monitor SARS-CoV-2 growth in cell culture supernatants that does not necessitate RNA extraction, and is as accurate and sensitive as existing methods. In a proof-of-concept screen, we found that E64D, apilimod, EIPA and remdesivir can substantially impede SARS-Cov-2 replication providing novel insight into viral entry and replication mechanisms. This facile approach will undoubtedly expedite basic SARS-CoV-2 research, be amenable to screening platforms to identify therapeutics against SARS-CoV-2 and can be adapted to numerous other RNA and DNA viruses.

## Observation

Severe acute respiratory syndrome coronavirus, SARS-CoV-2, the causative agent of the ongoing COVID-19 pandemic, is continuing to cause substantial morbidity and mortality around the globe (1, 2). Currently, there are no clinically approved countermeasures available for COVID-19 and the lack of a simple assay to monitor virus growth that can be used in basic SARS-CoV-2 research as well as drug screens is slowing progress in this area. Current Q-RT-PCR methods to quantify SARS-CoV-2 growth in cell culture supernatants rely on time-consuming and costly RNA extraction protocols (3). In this study, we developed a facile Q-RT-PCR assay that bypasses the RNA extraction steps, can detect viral RNA from as little as 5 μL of cell culture supernatants and works equally well with TaqMan and SYBR-Green-based detection methods.

A widely used assay to measure virus growth in the retrovirology field relies on determining the activity of reverse transcriptase enzyme from a small amount of cell culture supernatants (4), and we reasoned that we could adapt this approach to monitor SARS-CoV-2 growth. First, we tested whether the more stringent lysis conditions used to inactivate SARS-CoV-2 would interfere with the subsequent Q-RT-PCR step and affect the broad dynamic range obtained typically from purified RNAs. To do so, 5 μL of serially diluted SARS-CoV-2 N-specific RNA standards were mixed with 2x RNA lysis buffer (2% Triton X-100, 50 mM KCl, 100 mM TrisHCl pH 7.4, 40% glycerol, 0.4 U/μL of SuperaseIN (Life Technologies)), followed by addition of 90 μL of 1X core buffer (5 mM (NH_4_)_2_SO_4_, 20 mM KCl and 20 mM Tris–HCl pH 8.3). 8.5 μL of the diluted samples were each added in a reaction mix containing 10 μL of a 2x TaqMan RT-PCR mix, 0.5 μL of a 40x Taqman RT enzyme mix (containing ArrayScript™ UP Reverse Transcriptase, RNase Inhibitor), and 1 μL of primer/probe mix (2 μM Taqman Probe (/5’-FAM/TCAAGGAAC/ZEN/AACATTGCCAA/3’ Iowa Black FQ/) and 10 μM each of SARS-Cov-2 NC forward and reverse primers (5’-ATGCTGCAATCGTGCTACAA and 5’-GACTGCCGCCTCTGCTC)) in a final reaction volume of 20 μL. The reactions were ran using the following cycling parameters: 48 °C for 15 min, 95°C for 10 min, 50 cycles of 95°C for 15sec and 60°C for 1 min of signal acquisition. We found that the modified sample preparations did not impact the sensitivity, efficiency or the dynamic range of the Q-RT-PCR assay as evident in the virtually identical cycle threshold (Ct) values obtained for a given RNA concentration and the similar slopes of linear regression curves (Fig. 1A).

**Fig 1.**
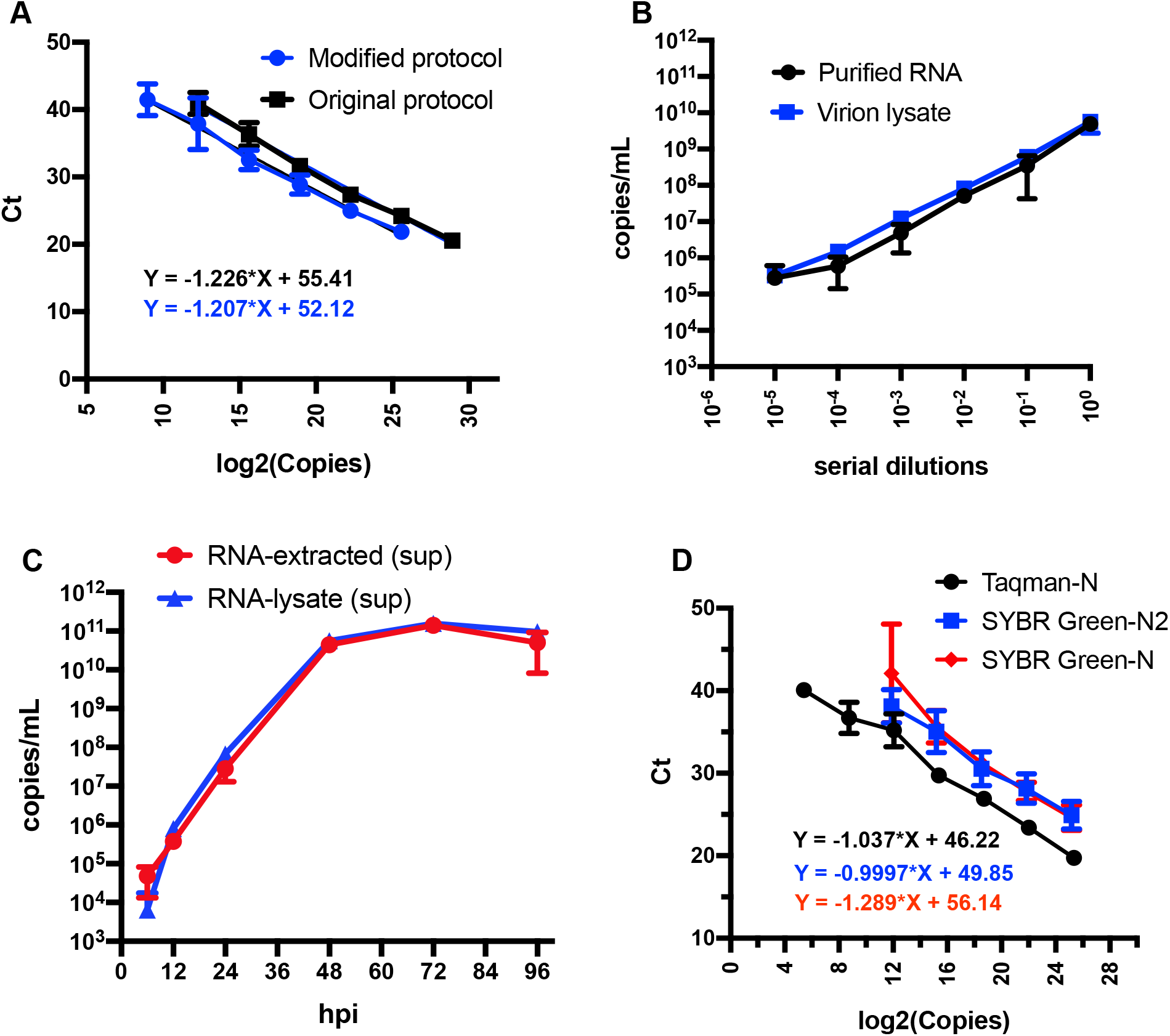
Development of a facile Q-RT-PCR assay for SARS-CoV-2 viral RNA detection in cell culture supernatants. (A) Serially diluted RNA standards were either directly subjected to Q-RT-PCR or processed as in the modified protocol detailed in the text prior to Q-RT-PCR. Log_2_ (copies) are plotted against the cycle threshold (Ct) values. Linear regression analysis was done to obtain the equations. Data show the average of three independent biological replicates. Error bars show the SEM. (B) Comparison of the efficiency and detection ranges for quantifying SARS-CoV-2 RNA using purified RNA or lysed supernatants from virus stocks. Data are derived from three independent replicates. Error bars show the SEM. (C) Vero E6 cells were infected at an MOI of 0.01 and cell culture supernatants were analyzed for SARS-CoV-2 RNA following the conventional RNA extraction protocol vs. the modified protocol developed herein at various times post infection. Cell-associated viral RNA was analyzed in parallel following RNA extraction for reference. Data are from three independent biological replicates. Error bars show the SEM. (D) Illustration of the efficiency and detection ranges of Taqman-based and SYBR-Green-based Q-RT-PCR quantifying known amounts of SARS-CoV-2 RNA. Data is from 2-3 independent replicates. Error bars show the SEM.

To determine whether this approach would work equally well for virus preparations, 100 μL of virus stock (1.4×10^5^ pfu) was lysed via the addition of an equal volume of buffer containing 40 mM TrisHCl, 300 mM NaCl, 10 mM MgCl2, 2% Triton X-100, 2 mM DTT, 0.4 U/μL SuperIN RNase Inhibitor, 0.2% NP-40. RNA was then extracted using the Zymo RNA clean and concentrator™-5 kit and was serially diluted afterwards. In parallel, 5 μL of virus stock and its serial dilutions prepared in cell culture media were lysed in 2X RNA lysis buffer and processed as above. Samples were analyzed by Q-RT-PCR alongside with RNA standards. We found that the modified assay performed equivalently well, if not better, with a similarly broad dynamic range (Fig. 1B).

We next used this assay to monitor virus growth on infected Vero cells. Supernatants containing virus collected at various times post infection were either used to extract viral RNA or subjected to Q-RT-PCR directly as above. The modified assay yielded virtually identical number of copies/mL of SARS-CoV-2 RNA in cell culture supernatants even at low concentrations of viral RNAs (Fig. 1C). Collectively, these results suggest that RNA extraction from cell culture supernatants can be bypassed without any compromise on the sensitivity or the dynamic range of Q-RT-PCR detection.

Next, we wanted to test whether this assay could work equally well with SYBR-Green-based detection methods. In addition to the N primer pair used in the above TaqMan-based assays, we utilized the N2 primer set designed by CDC and targeting the N region of the SARS-CoV-2 genome (F: 5’-TTACAAACATTGGCCGCAAA and R: 5’-GCGCGACATTCCGAAGAA). Serially diluted RNA standards were processed in RNA lysis and core buffers, and 7.5 uL of each dilution was used in a 20 uL SYBR-Green Q-RT-PCR reaction containing 10μL of a 2X POWERUP SYBR Green mix (Life Technologies ref: A25742), 1.25units/ μL of MultiScribe Reverse Transcriptase (Applied Biosystems), 1X random primers and 0.25 μM each of F and R primers. Both primer pairs yielded reasonably broad dynamic ranges, but were modestly less sensitive than TaqMan-based assays with a detection limit of ~3500 RNA copies/mL (Fig. 1D). In the following experiments, we decided to use the N2 primer set as it appeared to have a modestly enhanced sensitivity and efficiency overall (Fig. 1D).

One immediate application of this simplified assay is screening platforms given the ability to determine virus growth in small quantities of cell culture media. To demonstrate this, we next conducted a proof-of-concept drug screen to validate the antiviral activities of various compounds that have been reported to inhibit SARS-CoV-2 and HIV-1 replication as well as non-specific entry inhibitors (Table S1). Vero E6 cells plated in 96-well plates were infected in the presence of varying concentrations of the indicated compounds. Viral RNA in cell culture supernatants was quantified by the SYBR-Green-based Q-RT-PCR assay as above at 6, 24 and 48 hpi. Compound cytotoxicity was assessed in parallel by the RealTime-Glo™ MT Cell Viability Assay (Promega). While viral RNA was at background levels at 6 hpi (data not shown), we found that, at 24hpi, remdesivir (inhibitor of RNA-dependent RNA polymerase, (5)), E64D (inhibitor of the endosomal protease cathepsin B, K and L), and apilimod (PIKfyve inhibitor resulting in endosomal trafficking defects, (6, 7)) substantially decreased SARS-CoV-2 viral RNAs in supernatants (Fig. 2). IC_50_ values of these compounds (2.8 μg/mL (remdesivir), 3.3 μM (E64D) and 12nM (apilimod)) were within the same range of published IC_50_ values of these compounds (6–8) (Fig. 2). Similar results were obtained at 48 hpi, albeit E64D and apilimod appeared to be less potent at this time point either due to virus overgrowth or compound turnover (data not shown). We found that EIPA, which inhibits Na^+^/H^+^ exchanger and macropinocytosis, substantially decreased viral RNA in supernatants at sub-cytotoxic levels (Fig. 2D), suggesting that macropinocytosis may contribute to viral entry and/or subsequent steps in virus replication. HIV-1 specific inhibitors nevirapine, amprenavir and ALLINI-2 modestly inhibited SARS-CoV-2 replication without apparent cytotoxicity at high concentrations, albeit the concentrations required for this inhibition were much higher than those that inhibit HIV-1 (Fig. S1). Overall, these findings demonstrate that this miniaturized assay can be adapted for screening platforms and support previous reports which demonstrated that SARS-CoV-2 entry is dependent on processing of the Spike protein by cellular proteases and requires endosomal fusion (7, 9, 10).

**Fig 2.**
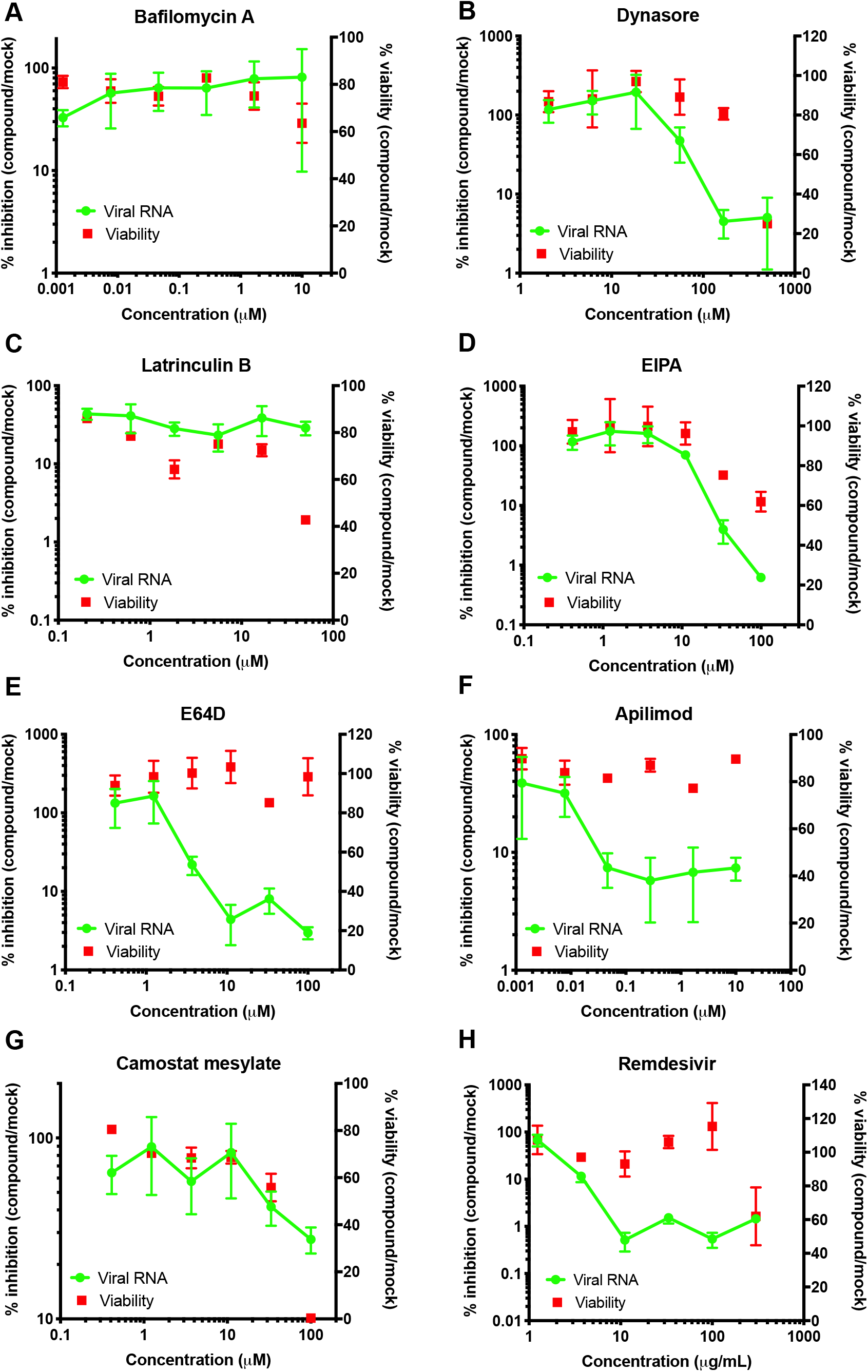
A compound screen to validate SARS-CoV-2-specific inhibitors and entry pathways. Vero E6 cells were infected with SARS-CoV-2 at an MOI of 0.01 and inhibitors were added concomitantly at concentrations shown in the figures following virus adsorption. Supernatants from infected cells were lysed and used in a SYBR-Green based Q-RT PCR to quantify the viral RNA in cell culture supernatants. Compound cytotoxicity was monitored by RealTime-Glo™ MT Cell Viability Assay Kit (Promega) in parallel plates. Data show the cumulative data from 2-5 independent biological replicates. Error bars show the SEM.

In conclusion, we have developed a facile Q-RT-PCR assay to monitor the kinetics of SARS-CoV-2 growth in cell culture supernatants bypassing the time consuming and costly RNA extraction procedures. This facile assay will undoubtedly expedite basic SARS-CoV-2 research, might be amenable to mid-to high-throughput screens to identify chemical inhibitors of SARS-CoV-2 and can be applicable to the study of numerous other RNA and DNA viruses.

## ACKNOWLEDGEMENTS

We thank members of the Diamond, Whelan, Boon and Amarasinghe labs for their generosity in providing reagents and support. This study was supported by Washington University startup funds to SBK, NIH grant U54AI150470 (the Center for HIV RNA Studies) to SBK, NSF grant DGE-1745038 to HRV and Stephen I. Morse fellowship to MPC.

---

**Fig S2.**
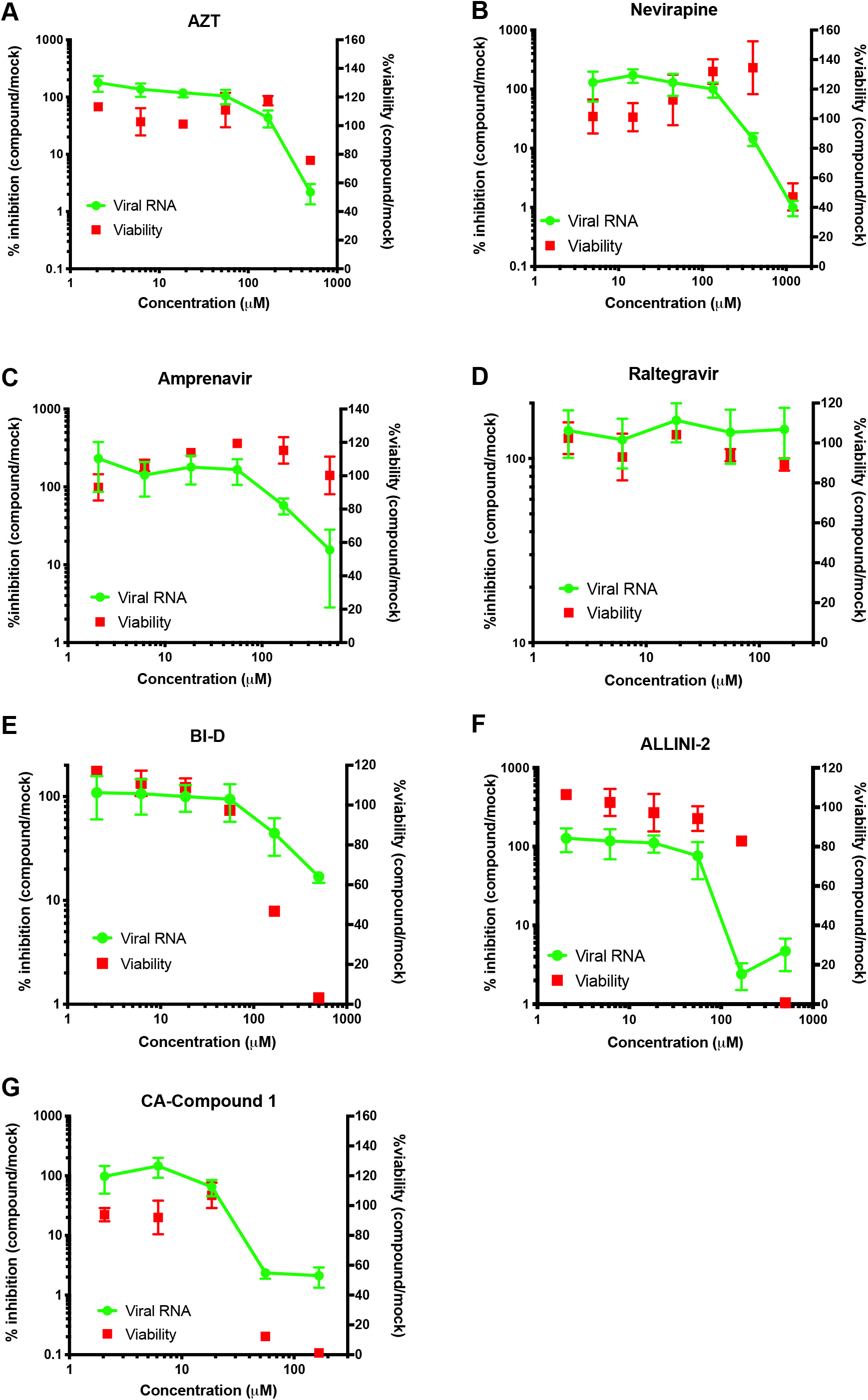
A screen to test the antiviral activities of various HIV-1-specific inhibitors. Vero E6 cells were infected with SARS-CoV-2 at an MOI of 0.01 and inhibitors were added concomitantly at concentrations shown in the figures following virus adsorption. Supernatants from infected cells were lysed and used in a SYBR-Green based Q-RT PCR to quantify the viral RNA in cell culture supernatants. Compound cytotoxicity was monitored by RealTime-Glo™ MT Cell Viability Assay Kit (Promega) in parallel plates. Data show the cumulative data from 2-3 independent biological replicates. Error bars show the SEM.

